# Extending of imaging volume in soft x-ray tomography

**DOI:** 10.1101/2022.05.11.491437

**Authors:** Axel Ekman, Jian-Hua Chen, Bieke Vanslembrouck, Carolyn A Larabell, Mark A Le Gros, Venera Weinhardt

## Abstract

Soft x-ray tomography offers rapid whole single cell imaging with a few tens of nanometers spatial resolution without fixation or labelling. At the moment, this technique is limited to 10 µm thick specimens, such that applications of soft x-ray tomography to large human cells or multicellular specimens are not possible. We have developed a theoretical and experimental framework for soft x-ray tomography to enable extension of imaging volume to 18 µm thick specimens. This approach, based on long depth of field and half-acquisition tomography, is easily applicable to existing full-rotation based microscopes. This opens applications for imaging of large human cells, which are often observed in cancer research and cell to cell interactions.

## 1. Introduction

Living organisms are composed of a variety of different cells. In humans alone, a rough estimate reveals more than 400 distinct cell types^1^. Understanding the internal organization of these cells is key for comprehensive structure-function analyses of cells and the varied responses to both external environment and diseases. Different cell types are also diverse in shape and size^2,3^, and it remains difficult to non-invasively image their entire internal structure with a single imaging technique. Among existing structural imaging modalities for cell biology, soft x-ray tomography (SXT) uniquely enables whole-cell imaging with a few tens of nanometers spatial resolution, minimal sample preparation, and quantitative contrast of protein-rich intracellular structures^4–10^. Due to these unique characteristics, SXT recently has been used for a wide range of applications, from phenotyping of cells such as bacteria, yeast, and advanced eukaryotes^11–16^ to investigating cell-nanoparticle^17,18^, cell-parasite^19–21^ and cell-virus interactions^22–27^.

SXT is suitable for imaging a wide variety of cell types; however, some investigators have reported that soft x-ray imaging is limited to specimens less than 10 µm thick^28–30^. Factors that affect the size of specimen include soft x-ray transmission through dense organic materials and the depth of field (DOF) and field of view (FOV) of the microscope and optics used.^31^ Cryopreservation methods, known to be important for cryoEM, are also of concern when preparing cells for SXT^29^. Better vitrification of thick cells has been addressed by modifying methods for plunge and high pressure freezing.^29,32,33^

Different approaches have been used to overcome the DOF limitations. Some investigators have chosen to work around the DOF issues by limiting the specimens they image to those less than 10 µm thick (at maximum tilt for limited-tilt tomography of extended “*flat”* specimens) or by imaging very small, thin regions of larger cells. Imaging with high-resolution optics (35nm optics) that have a shallow DOF further restricts the specimen size that can be imaged. Other investigators have demonstrated that it is possible to image larger specimens by using optics with a greater DOF but lower resolution (60- and 80-nm resolution zone plates) and by using full-rotation tomography. This latter approach was recently used to image whole pancreatic β cells and human lung epithelial cells, both larger than 10 µm^16,26^. There also have been efforts to extend the DOF of SXT without compromising resolution by using through-focus image acquisition schemes^34,35^ or by using modified reconstruction algorithms^31,36^, but these approaches have yet to be widely adopted.

The principles of image formation in SXT have been discussed in significant detail in previous publications^37–42^. In this manuscript, we discuss only those aspects that limit the size of cells that can be imaged with SXT. Based on the minimum soft x-ray transmission required for tomographic reconstructions and previously measured DOF and FOV of our full-field SXT microscope^36^, we suggest that imaging cells thicker than 18µm is feasible. Guided by these considerations, we adopted a “half-acquisition” tomographic data collection strategy to overcome the limited FOV at high spatial resolution (35 nm). We also employed a broad-band illumination (ΔE/E∼0.01) to overcome DOF limitations when using a 35 nm resolution zone plate objective lens. Using this approach, we demonstrated that SXT is capable of tomographic imaging, at high resolution, whole cells twice the size previously reported.

## 2. Theoretical Considerations of SXT Specimen Size Limitations

X-rays in the soft (low) energy range, particularly from 284 eV to 543 eV (λ = 2.34 nm to 4.4 nm), are superior in terms of contrast for imaging cells in their native, fully hydrated state. The beneficial contrast of soft x-rays in this so-called “water window” (between the x-ray absorption K-edge of carbon, at 284eV, and oxygen, at 543 eV) is demonstrated by plotting the attenuation length of water, protein, lipid droplets, and carbohydrates (**Figure 1a**). In the water window, carbon- and nitrogen-rich cellular structures of micrometer size attenuate x-rays 10 times more than water. The attenuation length is the thickness of a material where the flux of transmitted x-rays has dropped to (1/e) of the initial flux; this is equivalent to 63 % of the incoming x-ray photons being absorbed^46^. The attenuation length of water is of the order of 10 µm at water window imaging wavelengths. Biological cells are comprised mostly of water, ∼70 % by weight, and to date most biological imaging using SXT has been performed on cells about 10 µm or smaller^28–30,43–45^. The contrast in a transmission imaging technique such as SXT comes from transmitted x-rays. In the case of 10 µm-thick fully hydrated cell, there are still 37 % of the photons available for image formation. Due to the broad applications of x-ray absorption-contrast computed tomography, there are medical and industrial standards (EN16016-2, ISO 15708-2) stating that the minimal transmission required for obtaining an adequate tomographic reconstruction is 10-20 %. In some specific materials, optimal transmission of ∼10 %^47 48^ has been reported. Applying 10 % minimal transmission to SXT, water samples thicker than 20 µm can be imaged (**Figure 1b**). In addition, the increasing computing power available today enables iterative or deep learning reconstruction algorithms that are optimized for low signal-to-noise ratio^49–52^, this might make SXT imaging with as little as 5 % x-ray transmission feasible.

**Figure 1.**
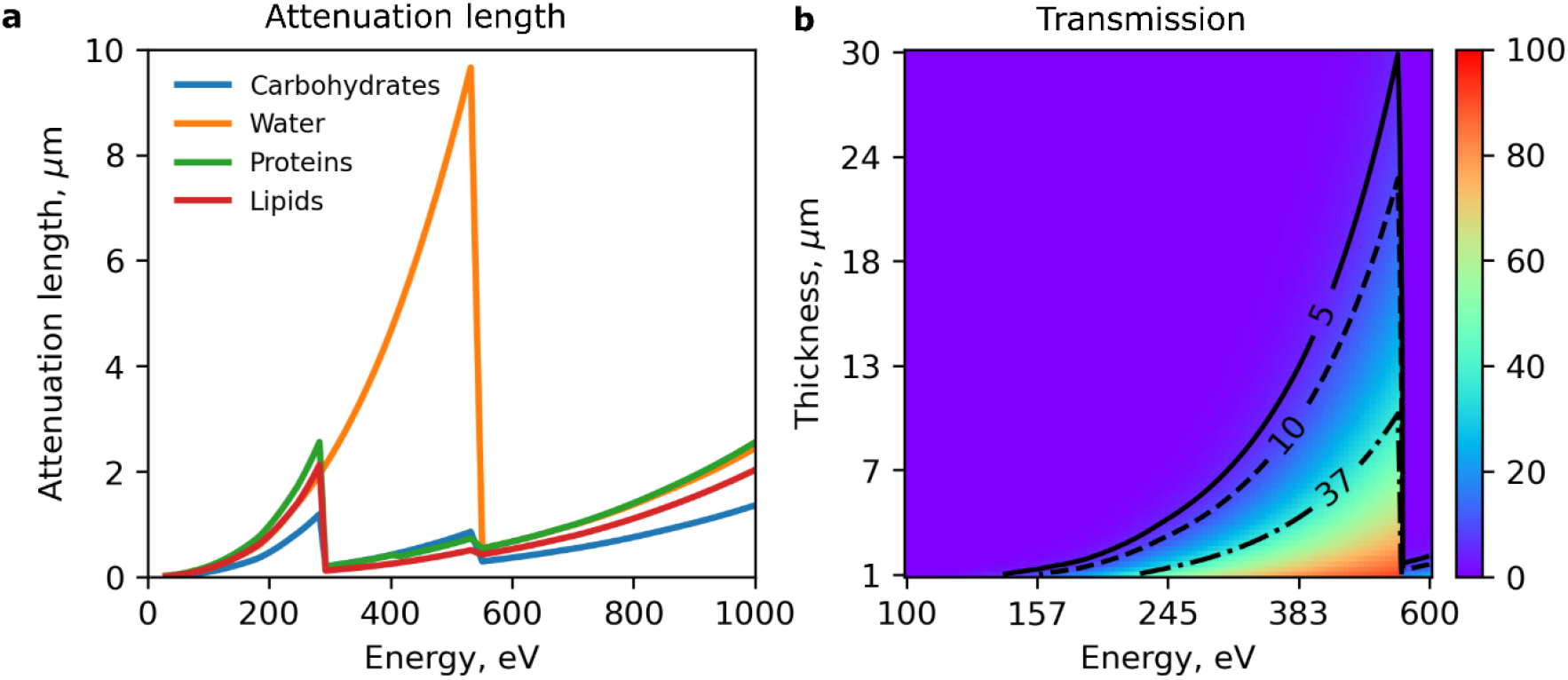
Attenuation length and x-ray transmission of organic materials in the soft x-ray energy range. a) attenuation length of typical carbohydrates (C_2_H_2_O_2_, 2.2 g/cm^3^), proteins (C_22_H_10_N_2_O_5_, 1.43 g/cm^3^), lipids (C_16_H_32_O_2_, 2.2 g/cm^3^) and water (H_2_O, 1 g/cm^3^). The absorption of carbon-rich materials dominates in the “water window” of x-rays, i.e. from 284 eV to 543 eV (2.34 nm to 4.4 nm). b) soft x-ray transmission in % with varying water thickness. Transmission of 37 % corresponds to the attenuation length of water in the “water window” and corresponds to a thickness of 10 µm. 10 % and 5 % x-ray transmission can be achieved in water of 23 µm and 30 µm respectively.

To obtain the highest quality 3D reconstruction with SXT, the whole specimen should remain in focus in each projection image. Because of this, the DOF of the microscope optics is one factor that limits imaging larger cells. Similar to visible light microscopy, the DOF in SXT depends on the numerical aperture (NA) of the objective lens, with 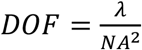. The objective lens of a soft x-ray microscope is typically a circular diffraction grating also called a Fresnel zone plate (FZP), and the DOF for monochromatic illumination at water window imaging wavelengths (∼2.4 nm) is just a few micrometers^5,31^ (**Figure 2a**). Thus, even for cells smaller than 10 µm, several studies have been dedicated to mitigating the limited DOF by collecting additional tomographic data at different focal positions^34,35,53^ or by including the DOF into the reconstruction algorithm^35,36^. Generally, such approaches lead to 3 times increase in the DOF (**Figure 2a**). Previous work demonstrated that in reality the DOF is actually elongated due to the existence of partial transverse coherence, quasi-monochromatic illumination, imperfections in Fresnel zone plates, and alignment of the optical components in the beam path^31,54,55^. Indeed, the experimentally measured DOF in the SXT microscope (XM-2) at the Advanced Light Source of the Lawrence Berkeley National Laboratory is 18 µm using a 60 nm zone plate at 2.4 nm wavelength^36^. Clearly, such an extended DOF allows imaging larger cells.

**Figure 2.**
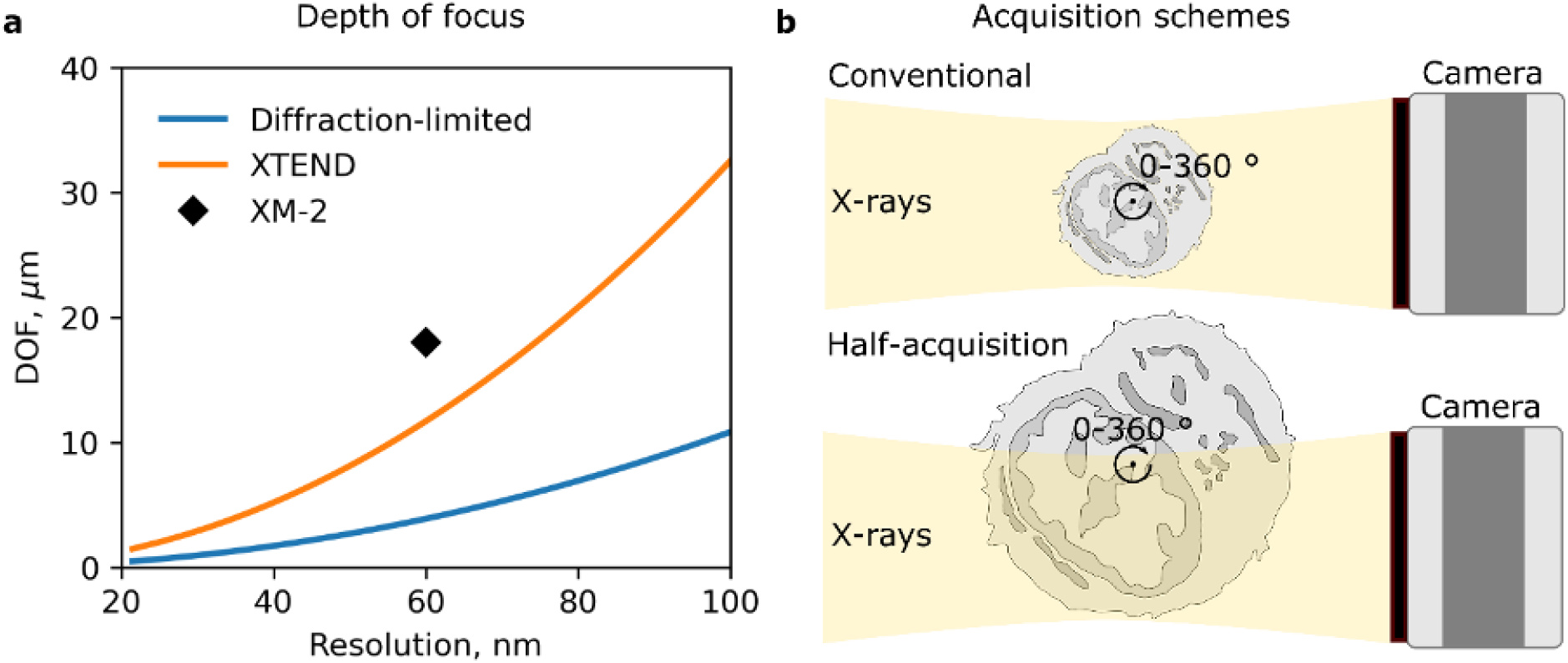
Achievable depth of field and field of view in soft x-ray tomography. a) diffraction-limited, computationally extended (XTEND^35^) and experimental (from XM-2^36^) depth of field in SXT microscopes as a function of spatial resolution. b) schematics of conventional (top) and half-acquisition(bottom) schemes for x-ray tomography. Lateral increase in field of view in half acquisition is achieved with full rotation (0-360°) tomography.

In addition to sufficient x-ray transmission and extended DOF, imaging larger specimens requires matching the field of view to the cell size. As SXT aims to provide high spatial resolution, the pixel size on the camera after magnification is on the order of 10 nm. Therefore, with a typical sensor size of 1Mpixels, a FOV up to 10 µm can be imaged in a single exposure. All imaging microscopes have a finite FOV, and a variety of approaches have been used to image specimens larger than the FOV. In commercial computed tomography (CT) scanners, as well as laboratory and synchrotron-based hard x-ray tomography instruments, “wide-field” or “half-acquisition” tomography has been used to expand the lateral FOV^56–59^. For instruments capable of full rotation data collection, half-acquisition methods are superior to the more traditional tiled acquisition approach. This is because the specimen only requires rotation to collect tomographic data, and translation normal to the optical axis is not required.

The half-acquisition approach is based on the symmetry of tomographic data acquisition, where the specimen is rotated around its central axis, see **Figure 2b**. If the specimen fits within the FOV, rotation though 180 degrees is sufficient to collect a complete isotropic resolution dataset (conventional case, **Figure 2b top**). At each rotation angle, x-ray projection provides a line integral along the x-ray path, such that the projections at 0 and 180 degrees are identical and a complete 3D volume can be reconstructed from half rotation of the specimen. For a specimen larger than the FOV, the rotation axis can be shifted to lie along the edge of the imaging CCD. If data is collected through a full (360 degree) rotation of the specimen, then a complete data set has been collected (half-acquisition case, **Figure 2b bottom**). The symmetry of the collection method has then been used to effectively double the FOV. Until now, the half-acquisition scheme has been applied only with parallel and cone x-ray beam geometries.

In total the preceding discussion of the transmission of soft x-rays through biological material, the experimental DOF, and the extension of the FOV via the half-acquisition data collection scheme indicates that SXT imaging of cells up to ∼20 µm in diameter should be possible.

## 3. Experimental Result

The experiments were performed on the soft x-ray microscope (XM-2) at the Advanced Light Source of the Lawrence Berkeley National Laboratory^60^. The microscope is equipped with a sample stage that allows full 360 degrees specimen rotation and two “switchable” FZP objective lenses^61^, enabling 3D imaging with 35 nm and 50 nm spatial resolution. We first used the half-acquisition data collection geometry for SXT imaging on human B cells. These cells match the previously reported specimen thickness of 10 µm. To exploit the effective increase in sensor size with the half-acquisition protocol, we used the high resolution FZP with 35 nm spatial resolution. **Figure 3a** shows a pair of corresponding mirror projections (0 and 180 degrees) prior to merging (top), and after merging (bottom) using weighted averaging (see methods). The merged x-ray projections were then reconstructed using the SXT reconstruction software^62^ into a 3D volume (**Figure 3b and c**). By imaging using the half-acquisition geometry, we obtained an SXT volume with an effective FOV of 14 µm x 14 µm (instead of the actual 9 µm x 9 µm FOV dictated by the sensor size and microscope magnification) using the 35nm resolution FZP. The increase in the FOV is less than anticipated, due to the need to overlap the two regions and the need to account for a slight specimen shift during rotation, see **Video 1**. In some datasets not all mirror projections had sufficient overlap, resulting in “missing information” artifacts in the reconstructions, see **Video 2**. Additionally, in all (n=5) datasets, organelles on the periphery of the reconstructed volume appeared doubled (**Video 2**), indicating that the DOF of the 35 nm FZP is less than 14 µm. Nevertheless, the half-acquisition scheme can be used to visualize fine organelle structures, for example ER and mitochondria contacts as shown in **Figure 3c**, at high spatial resolution without a decrease in the FOV.

**Figure 3.**
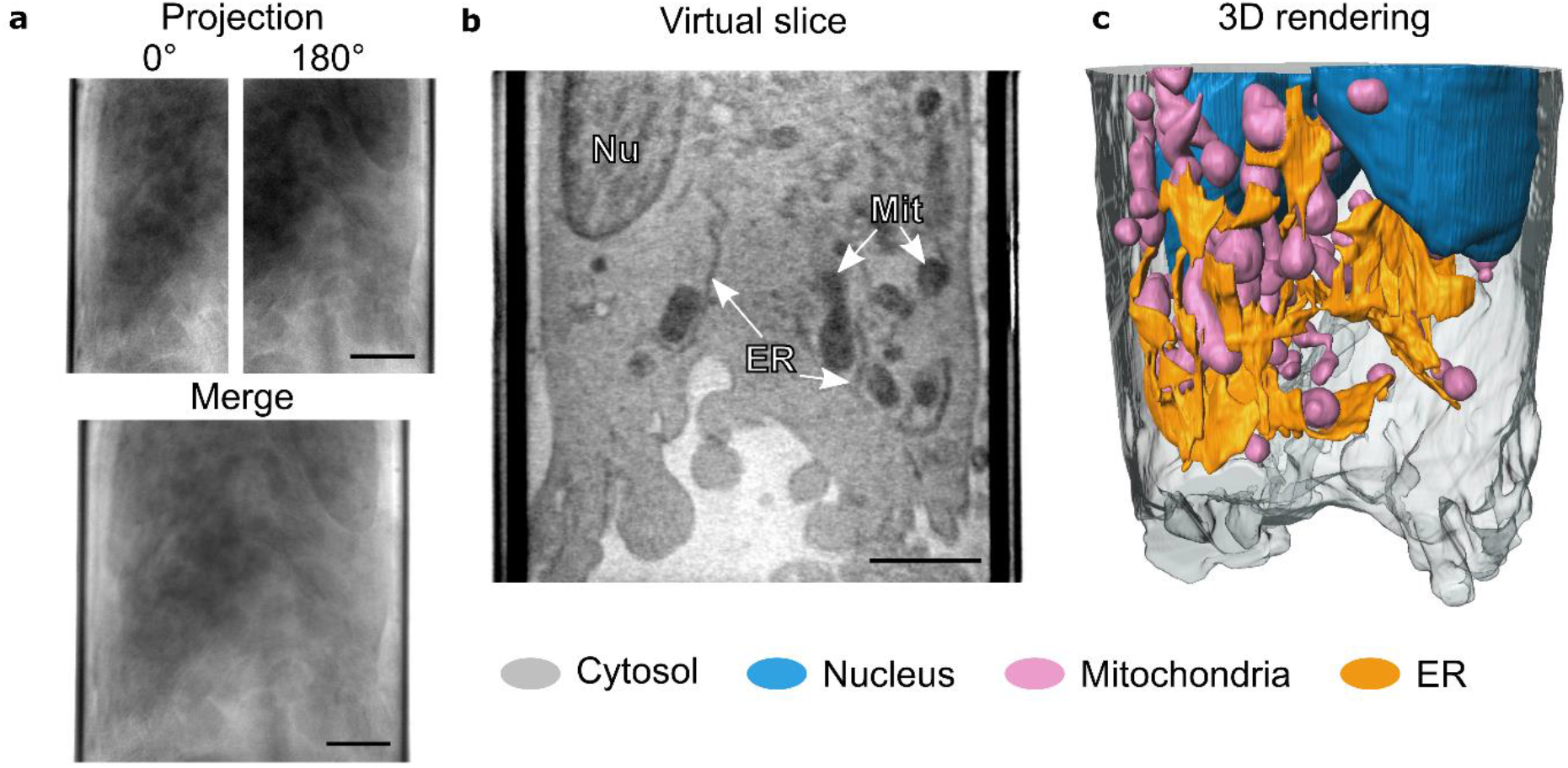
Extension of the field of view in SXT imaging via the half-acquisition scheme using a 35 nm FZP. a) x-ray projections at 0° and 180° rotation angles before and after merging, b) reconstructed virtual slice showing the nucleus (Nu), mitochondria (Mit), and endoplasmic reticulum (ER). The LAC is scaled from 0.1-0.6 pm^-1^, c) 3D rendering of a cell acquired using the half-acquisition scheme with a 35 nm FZP, showing segmented organelles. The scale bars are 2 μm.

We then explored the use of the half-acquisition data collection scheme using a 50 nm FZP objective to image human osteosarcoma epithelial cells (U2OS), which are typically about 20 µm diameter^63^. We obtained sufficient x-ray transmission in 16-18 µm thick specimens, some of which contained multiple cells that were densely packed with organelles (**Figure 4a**). We did not observe any artifacts, such as streaks or additional noise, in these large SXT volumes, see **Video 3**. Blurring of the organelles was not visible with the 50 nm FZP, which validates the previously measured extended DOF at the XM-2 microscope (**Figure 2a**). The LAC of analyzed organelles in these tomograms was identical to those previously reported^26^. **Figure 4b** shows a 3D rendering of an 18 µm x 18 µm x 13 µm volume acquired using SXT. The experimental data validates the use of a 50 nm FZP combined with our half-acquisition collection protocol to image specimens thicker than 18 µm.

**Figure 4.**
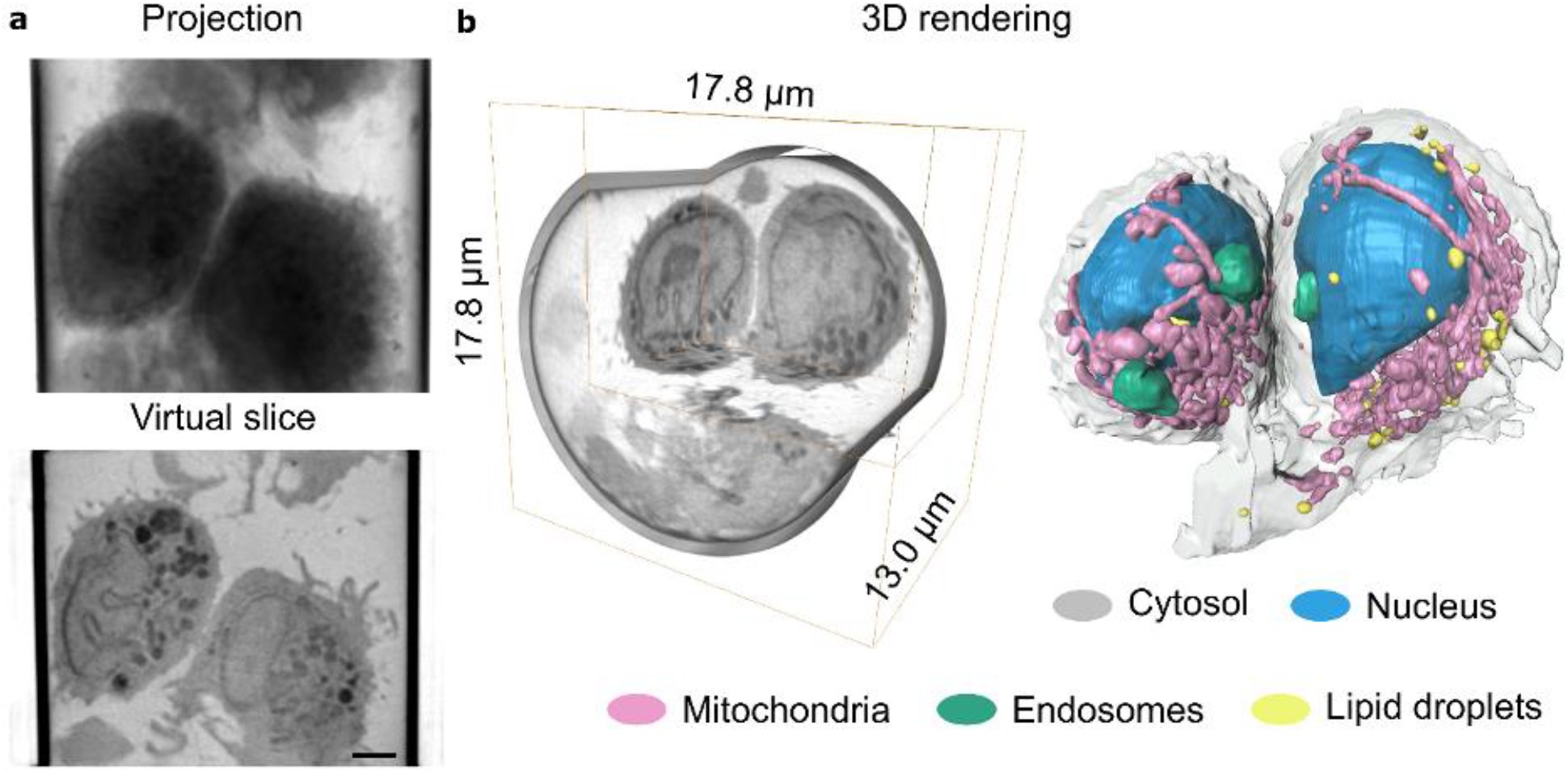
Extension of the SXT imaging volume using the half-acquisition scheme with a 50 nm FZP. a) merged mirroring projections (top) and virtual slice through the reconstructed SXT volume (bottom) of two U2OS cells inside a glass capillary, b) 3D rendering of the SXT volume (left) and segmentation of cells (right) shown in a. SXT volume of 17.8 μm × 17.8 μm × 13.0 pm comprises two organelle-rich U2OS cells, ice, and surrounding dense glass capillary. The LAC is scaled from 0.1-0.6 μm^−1^. Scale bars are 2 μm.

## 4. Conclusion

We have outlined methods and presented experimental results that extend the previously reported limit of penetration depth in SXT cellular imaging from 10 µm to 18 µm. To increase the specimen volume imaged by SXT, we combined the long DOF performance of a broadband-illumination x-ray microscope with a half-acquisition data collection scheme applicable to a cylindrical specimen geometry. These methods enable a 4X increase in imaging volume and are most suited to synchrotron-based instruments with full rotation data collection capability such as the XM-2 microscope at the Advanced Light Source.

For the XM2 microscope, we anticipate further improvements in data collection with future optimization of the sample tomographic stage to correct for imperfections in the mechanical rotation axis. Future implementation of through-focus methods combined with half-acquisition data collection strategies will also allow even better resolution large volume datasets to be collected.

Given the increasing importance of SXT as a whole cell imaging method the ability to increase the imaging volume, opens new SXT applications for imaging of large human cells, which are typical for cancer, cell to cell interactions and even multicellular tissue.

## 5. Experimental Methods

### Analysis of soft x-ray transmission and depth of field

To calculate x-ray transmission, we have used tabulated refractive indexes for different compounds^46^. In our work we considered water (H_2_O, 1 g/cm^3^), carbohydrates (C_2_H_2_O_2_, 2.2 g/cm^3^), proteins (C_22_H_10_N_2_O_5_, 1.43 g/cm^3^) and lipids (C_16_H_32_O_2_, 2.2 g/cm^3^). Based on the Beer-Lamber law, x-ray transmission *T* decreases exponentially as a function: 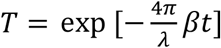. Here *λ* – is x-ray wavelength, *β* – is the imaginary part of the x-ray refractive λ index and *t* – is a thickness of compound. The attenuation length of compound is a specific case when transmission of the x-ray beam has decreased to a value of 1/e. The diffraction limited depth of focus (DOF) for x-ray Fresnel Zone Plates with a numerical aperture *NA*, is 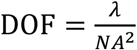. Based on Rayleigh criterion the diffraction-limited spatial resolution of a perfect lens is 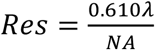. These basic simplified parameters have been used to demonstrate attenuation length and x-ray transmission with respect to x-ray energy, and DOF with respect to spatial resolution chosen for the SXT microscope.

### Cell culture

The U2OS cells were purchased from ATCC (HTB-96). The U2OS cells were maintained in Dulbecco’s Modified Eagle Medium (Invitrogen, Waltham, MA) supplemented with 10 % fetal bovine serum (Invitrogen) and 100 units/ml penicillin-streptomyocin (15140-122, Life Technologies, Carlsbad, CA).

The human B lymphocytes (GM12878) were purchased from the NGIMS Human Genetics Cell Repository, Cornell Institute of Medical Research (Camden, NJ). The B cells were maintained in Advanced RPMI-1640 medium supplemented with 15 % of fetal bovine serum, L-glutamine, (Gibco™, Carlsbad, CA) as a suspension.

Both cell lines were kept at 37 °C and 5 % CO_2_. The medium was refreshed 2-3 days per week to maintain a cell density of 0.2-1×10^6^ ml^-1^.

### Freezing of cells

The U2OS cells were detached by incubation in 0.25 % Trypsin for 5 min at 37 °C and 5 % CO_2_. Both cell types were collected by centrifugation at 125 x g for 10 min. The cells were loaded into thin-wall glass capillaries with a micro-loader. The capillaries were rapidly plunged into a liquid nitrogen cooled liquid propane and stored in liquid nitrogen until image acquisition^64^.

### Image acquisition

Capillaries were kept in a stream of liquid nitrogen-cooled helium gas during data collection^60,66^. To employ half-acquisition scheme, tomographic rotation was first aligned with respect to the center of the camera’s sensor and then shifter laterally “off-set” by half of the field of view. The x-ray projections were acquired by sequentially rotating capillaries with a 2 degrees angle to obtain a full 360°rotation. For normalization of the data, the series of 20 reference images were taken without capillaries with the same exposure time as for x-ray projections. The effective pixel size was 27 nm and 18.6 nm for the 50 nm and 35 nm Fresnel Zone Plates (FZPs) respectively. Depending on the thickness of a specimen and FZP used, the x-ray images were acquired with an exposure time of 200-300 ms.

### Reconstruction of half-acquisition dataset

In principle, the reconstruction can be calculated directly from the half-acquisition (HA) tomograms. By alignment of the projections, the projection operator (and it’s adjoint operator) can be constructed so that the geometry of the half acquisition is included in the projection matrix, and thus the reconstruction can be iteratively solved for. We opt here to artificially produce a complete sinogram, so that there is no need for specialized software and the dataset can be reconstructed by any standard reconstruction scheme.

The formation of the dataset for HA consists of choosing the corresponding ‘mirror’ pair from the tomographic data, inverting the horizontal axis, and registering the two images to form a complete, extended radiograph. Here, the images were aligned and combined using linear blending using the distance from the image border as the weights for the interpolation. The data was then aligned and reconstructed with AREC3D^62^.

### Alignment of the HA images

Due to the problem of partial overlap, the cost function of the minimization is not straightforward^67^. In **Figure 5** we show two examples of problems that might occur. One is that there may be multiple local maxima in the codomain of the function, making optimization difficult and often depending on the initial guess. The second one is that when the overlap between the images becomes small, it is uncertain whether a good alignment exists between the two images within the available search space.

**Figure 1a.**
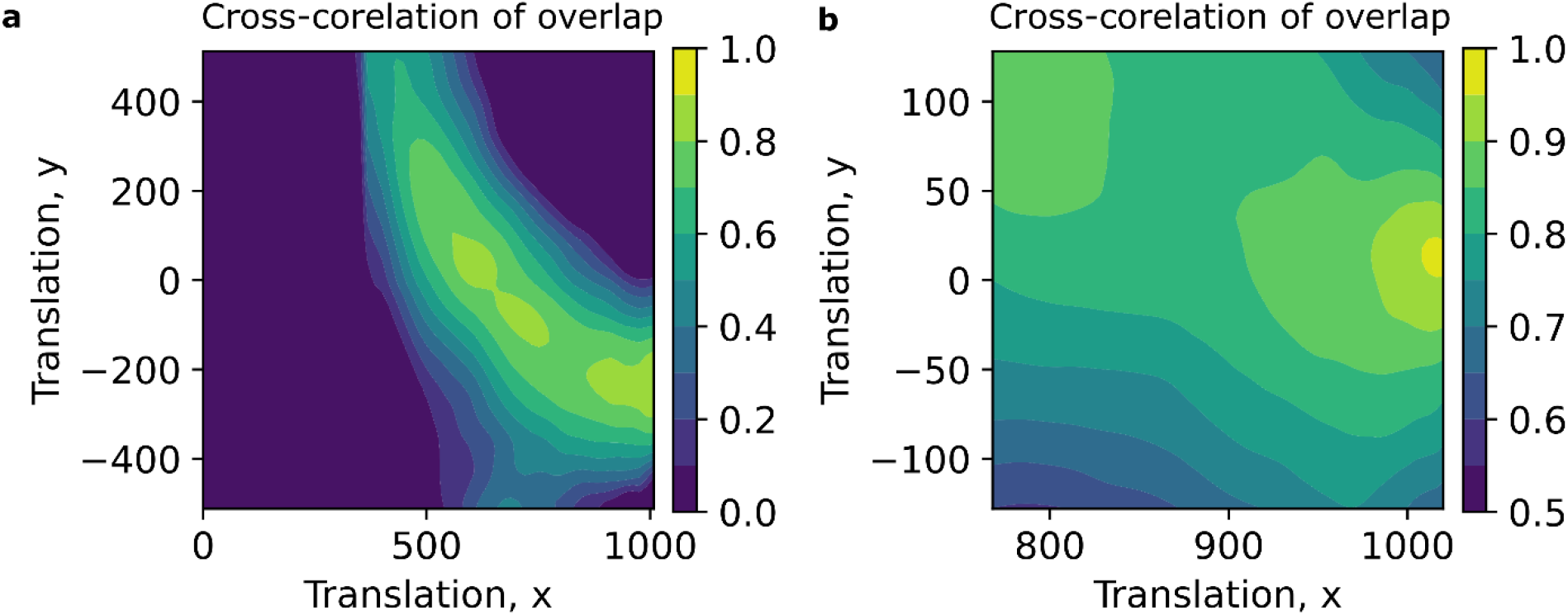
Two examples of the normalized cross-correlation of the overlapping area between a radiograph an its corresponding mirror image. a) Two local maxima of similar quality occur, without prior knowledge it is hard to determine which is correct. b) There is not sufficient overlap between the images to determine if a good alignment exists between the images.

### Initial guess

To address the problem of local maxima we want to build a robust initial guess for the alignment. For this we utilize that our samples are enclosed in a capillary tube, relatively cylindrical in shape, thus the width of the capillary tube is of similar width in each image.

The walls of the capillary are detected by standard edge-detection^68,69^. We used a horizontal Canny filter^70^ to detect the slopes, the locations were then defined for each row in the image as the location of the maximum of the filtered row. The location of the edge for each image *e*_*i*_ was defined as the median location of this edge.

For example, if the camera is focused on the left part of the sample, the width becomes

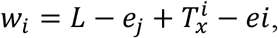

where *L* is the width of the image, 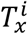 is the *x* component of the translation vector for image *i. e*_*i*_ and *e*_*j*_ are the edge location of the *i*th image and its corresponding mirror index *j*.

The prealignment of the capillary width is shown in Algorithm 1. A randomly chosen subset of all images are chosen for each iteration of the capillary alignment. The alignment is repeated until reaching a tolerance below 1 pixel.

#### Algorithm 1

Prealignment of the x-translation

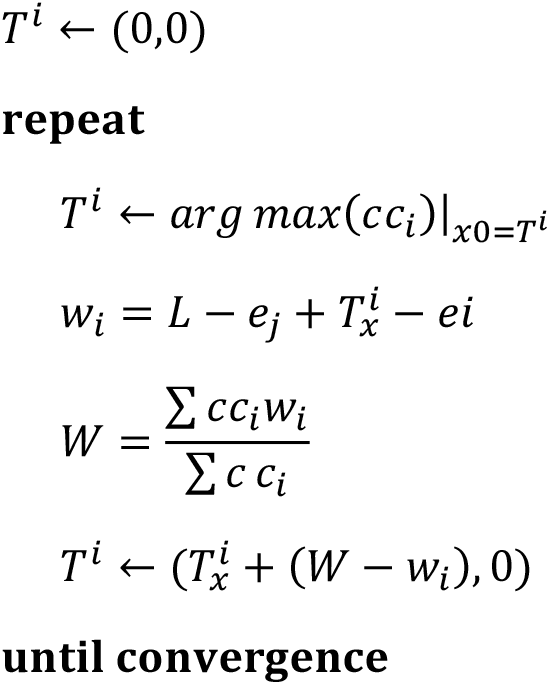

### Robust registering via neighbours

For the situations shown in **Figure 5b** we utilize the fact that successive images can be fairly accurately aligned. When direct alignment of image *i* to its mirror pair *j* is unstable or missing, an alignment can be found through its neighbouring images *i*–1 and *j*–1. A similar path though alignment graph can be constructed via successive alignments considering n nearest neighbours, see **Figure 6**.

Due to drift between successive image registrations, the obtained alignment from the *n* nearest neighbours is not well defined by a mean measurement of these paths, thus the final shift was obtained by a B-spline representation^71^ of the alignment parameter with respect to the distance. Error estimates were derived based on the deviation from the capillary width and the local variance in the shift estimate. The data points of the paths were weighted by the product of the geometric mean of the cross correlations in the path, the width of the overlap between the mirror correlation pair, and the inverse of the error estimate.

**Figure 2a.**
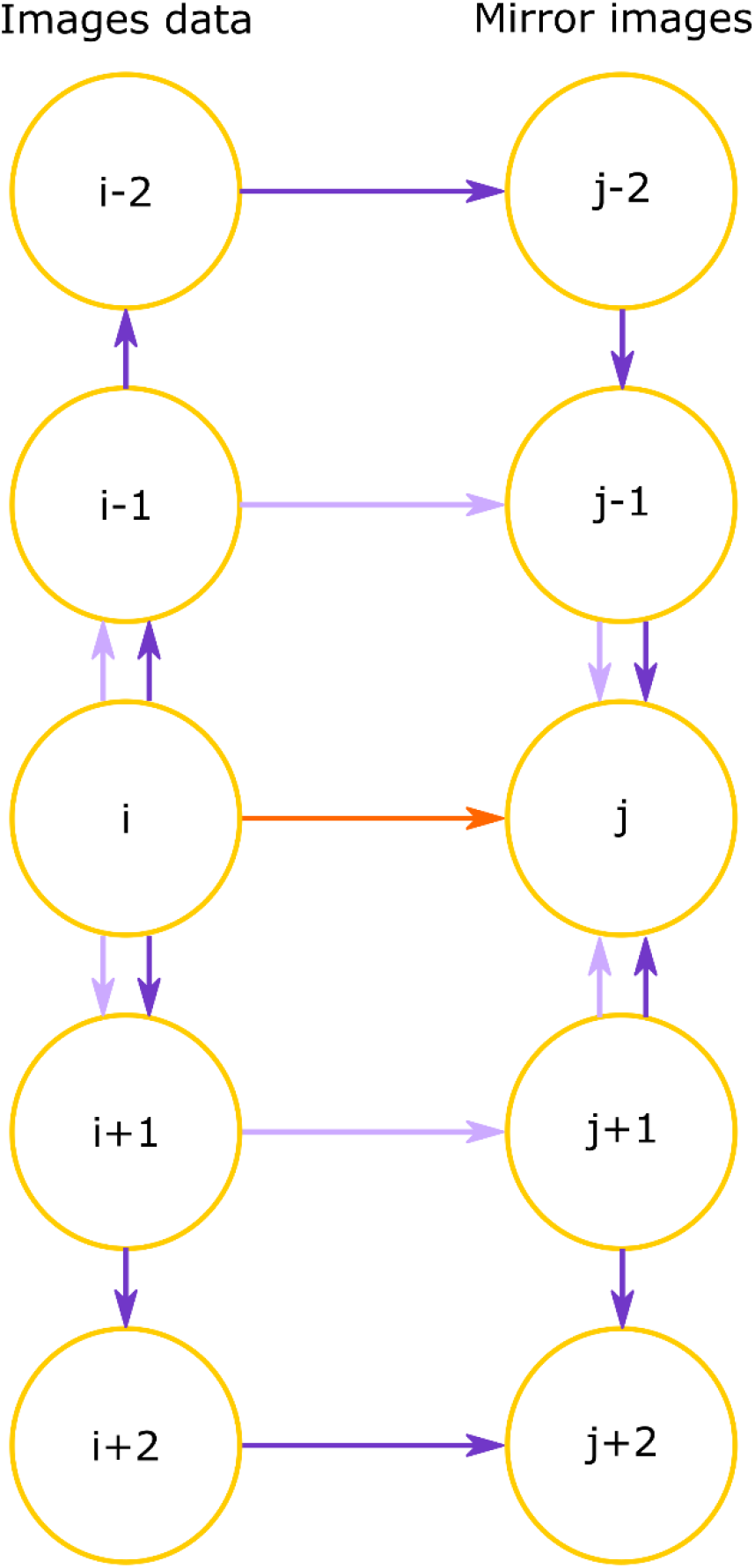
When direct alignment of image *i* to its mirror pair *j* is unstable or missing, an alignment can be found through its neighboring images.

### Segmentation and visualization

All cells were manually segmented using the software Amira 2019.1. The organelles were identified based on previously reported shape and linear absorption coefficient (LAC) values. Plots were prepared with Jupyter notebook^72^. Figures were assembled with Inkscape^73^.

## Video Legends

**Video 1**. SXT projection images acquired in half-acquisition mode with a 35 nm FZP. With an effective pixel size of 9.35 nm, precision of the rotation stage is insufficient to keep one-half of the sample in the field of view at all angles. Scale bars are 2 µm.

**Video 2**. Virtual slices through the reconstructed half-acquisition SXT dataset collected using a 35 nm FZP. SXT projections of the Cell 2 dataset could be merged at all angles, resulting in artifact-free virtual slices. The overlap of SXT projections for Cell 3 was insufficient at some angles, resulting in line artifacts of “missing information.” In both datasets, cellular structures in the first and last virtual slices, that is the periphery of the 3D volumes, are doubled. Scale bars are 2 µm.

**Video 3**. Virtual slices through the reconstructed half-acquisition SXT dataset with a 50nm FZP. The SXT volume comprises 17.8 µm x 17.8 µm x 13.0 µm of dense glass capillary, ice and two organelle-rich osteosarcoma cells (U2OS). The organelles are depicted as: cytosol – grey, nucleus – blue, mitochondria – pink, endosomes – green, and lipid droplets – yellow.

## Acknowledgements

V.W. is supported by German Research Foundation research fellowship WE 6221/2-1. The National Center for X-ray Tomography is supported by NIH NIGMS (P41GM103445, P30GM138441) and the DOE’s Office of Biological and Environmental Research (DE-AC02-5CH11231). V.W. was supported by the ERC Synergy Grant IndiGene (no. 810172 to Jochen Wittbrodt). V.W is supported by the CoCID project (no. 101017116) funded within EU Research and Innovation Act.

